# Production and evaluation of fluorophore-doped polymer substrates to screen for plastic-degrading enzymes

**DOI:** 10.1101/2025.07.13.664576

**Authors:** Anton A. Stepnov, Brenna Norton-Baker, Esteban Lopez-Tavera, Ravindra R. Chowreddy, Vincent G. H. Eijsink, Gregg T. Beckham, Gustav Vaaje-Kolstad

**Affiliations:** Faculty of Chemistry, Biotechnology and Food Science, NMBU - Norwegian University of Life Sciences, Ås, 1433 Norway; Renewable Resources and Enabling Sciences Center, National Renewable Energy Laboratory, Golden, CO, 80401 USA; BOTTLE Consortium, Golden, CO, 80401 USA; Norner Research AS, Porsgrunn, NO-3920 Norway

## Abstract

Fast and sensitive analytical methods are the key to efficient screening of plastic-degrading enzymes. While liquid and gas chromatography with mass spectrometry detection offer great power, the cost of the equipment is high, and the methods may not be straightforward when studying crude environmental samples. Here, we present a streamlined and affordable approach to assess the enzymatic deconstruction of insoluble synthetic polymers by blending them with a fluorescent probe, rhodamine 6G, and we evaluate this screening method using poly(ethylene terephthalate) (PET) as a model material. Our results indicate that enzymatic depolymerization of the rhodamine-doped PET can be observed in a high-throughput fashion by following release of the fluorophore. The fluorescence data obtained during the hydrolysis of rhodamine-doped PET by 14 engineered PET hydrolases, produced with a robotic platform, correlated with the quantitative chromatographic analysis of PET degradation products. Remarkably, the use of the rhodamine-loaded PET substrate resulted in negligibly low background signals even when detecting PETase activity in crude cell lysates, suggesting suitability for screening of a wide variety of samples. Encouraged by these results, we next produced a selection of polyethylene- and nylon-based materials loaded with rhodamine 6G. While rapid leaching of fluorophore observed with nylon substrates limits the utility of the method for detecting nylonase activity, the rhodamine-loaded polyethylene showed promising performance in passive diffusion tests, indicating that this latter substrate may be used to screen for polyolefin-degrading enzymes.

## Introduction

Despite recent advancements in the discovery of plastic degrading enzymes [1], product analysis still represents a major bottleneck in the search for novel catalytic activities. The diverse chemistry of synthetic polymers complicates method development and often necessitates the use of multiple techniques to test a large variety of potential enzyme/substrate combinations. Having access to powerful analytical methods such as liquid chromatography combined with tandem mass spectrometry (LC-MS/MS) does not resolve this problem completely. One may still need to use several different separation and detection conditions to ensure that all types of potential degradation products can be identified and quantified. Clearly, rapid and affordable methods for assessing plastic deconstruction in a standardized manner for multiple polymer types are of great interest to the field. Indeed, many such methods are being discussed and developed, based on various chemical and physical principles [2-4].

One of the ways to streamline screening for plastic-active enzymes with multiple polymer types is using substrates blended with fluorescent or fluorogenic probes. The deconstruction of such blends can be, in principle, observed in a high-throughput manner by following the fluorescence of the released tracer compounds instead of directly detecting the degradation products. This concept has seen some application in the study of enzymatic depolymerization of various materials [5-8], including poly(lactic acid) (PLA), poly(ε-caprolactone) (PCL), poly(butylene succinate-*co*-adipate) (PBSA), polyurethane (PU) and poly(ethylene terephthalate) (PET). Taken together, these previous studies provide experimental evidence that the amount of the released fluorophore indeed correlates with the degree of polymer degradation determined by other methods, such as liquid chromatography, pH-stat titration, or quartz crystal microbalance with dissipation monitoring.

Despite previous studies confirming the overall validity of this screening approach in combination with several types of polymers (including PET), fluorophore-doped plastics are yet to become a well-recognized part of the enzyme characterization toolkit. In our view, the most significant limitation of the existing methods lies in the substrate preparation techniques. To the best of our knowledge, all fluorophore-loaded plastic substrates reported in the literature so far were produced by slow evaporation of polymer solutions in organic solvents, resulting in the formation of thin films. Despite its simplicity, this approach has limited scalability and is time consuming. Another complication lies in the chemical properties of some of the previously employed tracer compounds. Zumstein *et al*. and Liu *et al*. [6, 7] suggested using a di-ester derivative of fluorescein (fluorescein dilaurate; FDL), instead of the non-modified fluorophore, to slow down passive leaching of the probe from the polyesters into solution. Importantly, FDL released from the substrate upon enzymatic hydrolysis is not fluorescent and requires further *in situ* conversion to fluorescein (by the same enzymes) to be detected. Although Liu *et al*. [7] have convincingly demonstrated that FDL-doped PET films can be used to screen PETase libraries in a high-throughput manner, the fluorescence signal in such a system will not only depend on the capacity of the target enzymes to deconstruct PET, but also on their ability to hydrolyze FDL. Notably, the rates of these two reactions may not be correlated.

Here, we evaluate an alternative, scalable approach for producing fluorophore-loaded plastic substrates to facilitate enzyme screening. Our method is based on melt blending of target materials with the highly fluorescent water-soluble probe, rhodamine 6G, followed by extrusion and pelletization. The resulting rod-shaped pellets (length, 3.0 ± 0.4 mm; diameter, 2.4 ± 0.5 mm; *n* = 10), are uniform and straightforward to handle when setting up enzymatic reactions in microcentrifuge tubes or microtiter plates. We produced and tested a 200 g batch of rhodamine-loaded PET substrate, observing a correlation between the fluorescence in reaction supernatants and the amount of soluble degradation products during reactions with 14 PET hydrolases (PETases). Importantly, the negligibly low background signal resulting from the use of this PET substrate allowed us to detect polymer degradation not only in well-defined reactions with purified enzymes, but also in experiments with crude *E. coli* lysates. Motivated by these findings, we used the same approach to produce polyethylene- and nylon-based materials loaded with rhodamine 6G. Our results show that the fluorophore is poorly retained by both nylon 6 and nylon 6,6. Conversely, the very slow background leaching of the probe from the polyethylene-based blend may allow for its use to screen for polyolefin-degrading enzymes.

## Results

### Preparation of fluorophore-doped PET substrate

After considering multiple water-soluble fluorophores, rhodamine 6G (referred to as “probe” below) was selected to be used as a tracer compound based on its affordability [9] and a very high quantum yield (Φ ∼0.9 in water) [10]. A commercial, low-melting point PET resin (ball-milled CumaPET L04 100 granules; DuFor Resins BV; Zevenaar, the Netherlands) with a crystallinity of 33.3 ± 0.7 % (according to DSC analysis; **Table S1**) was utilized as the carrier plastic. The compounding process involved melt blending of the materials at 250 °C using a twin-screw extruder, followed by water cooling and pelletization (see **Fig. S1** for more details). Despite widespread use of rhodamine 6G, data on its thermal decomposition are scarce. There is an indication that the dye may start degrading at ∼220 °C [11], therefore a relatively high 1% w/w fluorophore load was used. The blending/extrusion process yielded ∼200 g of uniformly red PET pellets, with an average mass of 17.3 ± 3.0 mg (**Fig. 1**). The crystallinity of the blended PET was considerably lower (4.8 ± 0.9%; **Table S1**) compared to the source material (33.3 ± 0.7%). Thus, the resulting substrate should be well accessible to PET-degrading enzymes, which cannot handle high crystallinity [12].

**Figure 1.**
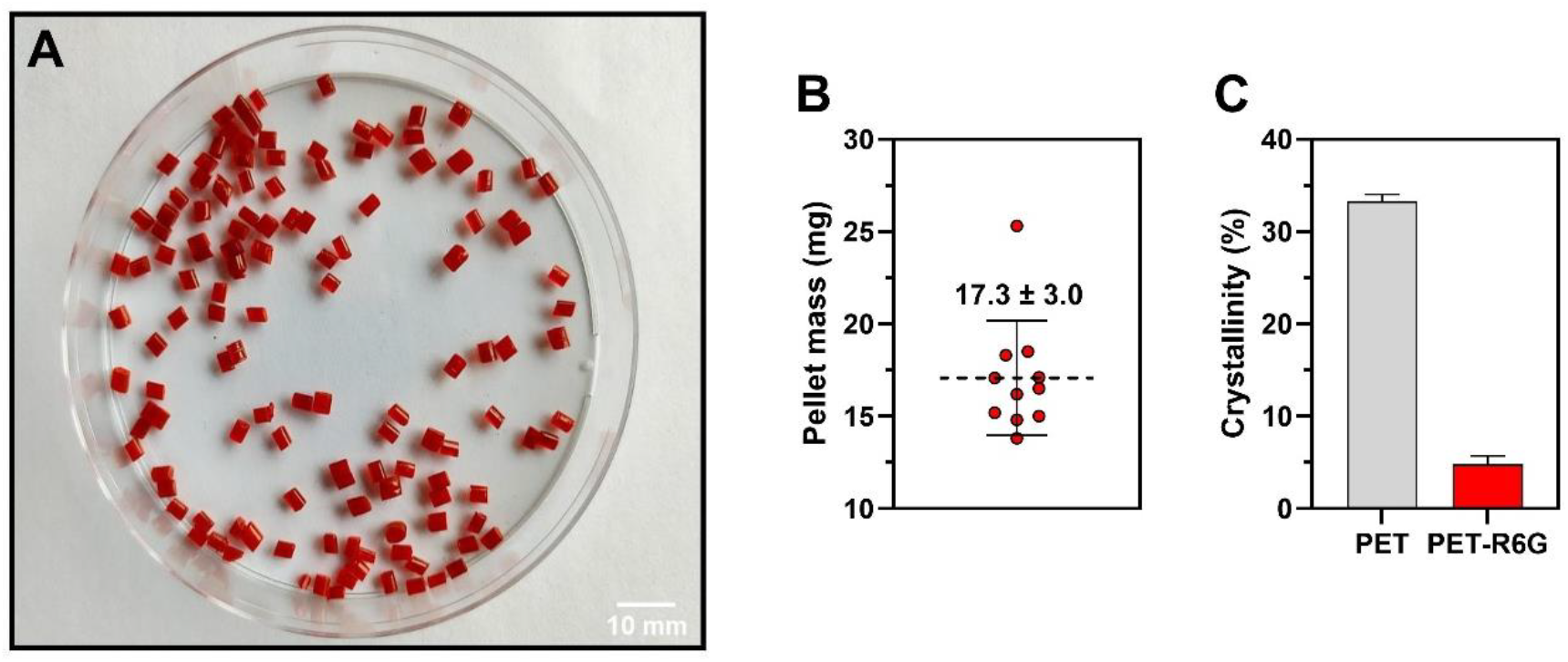
Fluorophore-doped PET (PET-R6G; A), the mass of individual pellets (B) and the crystallinity of the blended material. **(C)**. The error bars indicate the standard deviation between replicate measurements (**B**, *n*=11; **C**, *n*=2). Horizontal dashed line in **(B)** marks the average value. PET-R6G was produced by melt blending (250 °C) of milled PET granules with rhodamine 6G (1% w/w) using a twin-screw extruder with an L/D ratio of 25:1 operating at 500 RPM. The extrudate strands were water-cooled and then pelletized. See **Fig. S1** for a detailed description of the extrusion process. Source data are provided as a Source Data file.

Prior to performing proof-of-concept degradation experiments, the fluorescence of rhodamine 6G standard solutions, prepared in two different reaction buffers (sodium phosphate, pH 6.0, and Tris-HCl, pH 8.0), was measured. The relationship between the probe concentration and the fluorescence signal is increasingly non-linear at both pH values (**Fig. 2**) and can be well-described by a four-parameter logistic (4PL) model, commonly used to analyze fluorescence data [13].

**Figure 2.**
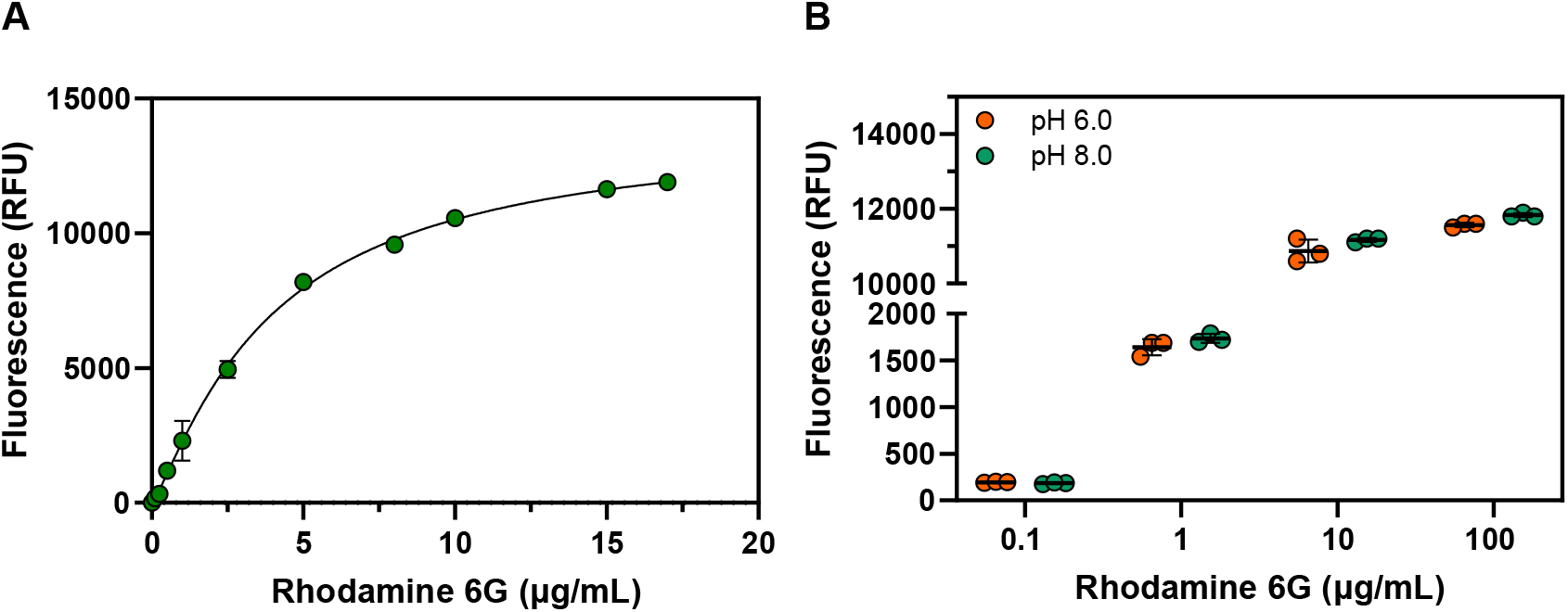
Fluorescence in standard solutions of rhodamine 6G. **(A)** The concentration dependence of rhodamine 6G fluorescence at pH 8.0 and **(B)** a comparison of rhodamine 6G fluorescence at pH 6.0 and 8.0 for various concentrations of fluorophore. Fluorescence was monitored at λ_ex/em_ = 530/552 nm in 50 mM sodium phosphate buffer (pH 6.0) or 50 mM Tris-HCl buffer (pH 8.0). Error bars indicate the standard deviation between triplicate measurements, whereas horizontal lines in **(B)** denote average values. The trend line in **(A)** represents a non-linear fit of the data using a four-parameter logistic (4PL) model (performed in GraphPad Prism v. 10.2.0). See the Source Data file for the regression parameters.

Notably, the 0-17 µg/mL fluorophore concentration range shown in **Fig. 2A**, approximately corresponds to 0-10% solubilization of a single 17 mg PET-R6G pellet in a 1 mL enzymatic reaction (assuming a precise 1% w/w rhodamine 6G load, homogeneous distribution of the fluorophore, and no adsorption of the released probe onto the reaction vessel inner walls). Determining higher concentrations of rhodamine 6G in reaction mixtures will require dilution due to significant deviation from 4PL model, likely caused by fluorescence quenching (**Fig. S2)**. On a side note, rhodamine 6G is a strong chromophore and can also be detected using absorbance spectroscopy at 530 nm (**Fig. S3**). Importantly, **Fig. 2** shows that the use of different buffer systems (sodium phosphate buffer/Tris-HCl) at different pH (6.0/8.0) does not affect the fluorescence signal (**Fig. 2B**).

### Probing enzymatic degradation of PET-R6G

Next, we tested whether rhodamine 6G is released upon the enzymatic hydrolysis of PET-R6G. Indeed, incubation of a single substrate pellet with a benchmark variant of the leaf branch compost cutinase (LCC-ICCG) [14, 15] resulted in fluorophore release that was linear over time for up to approximately 50 hours (∼0.2 µg/h; R^2^ = 0.98; **Fig. 3**).

**Figure 3.**
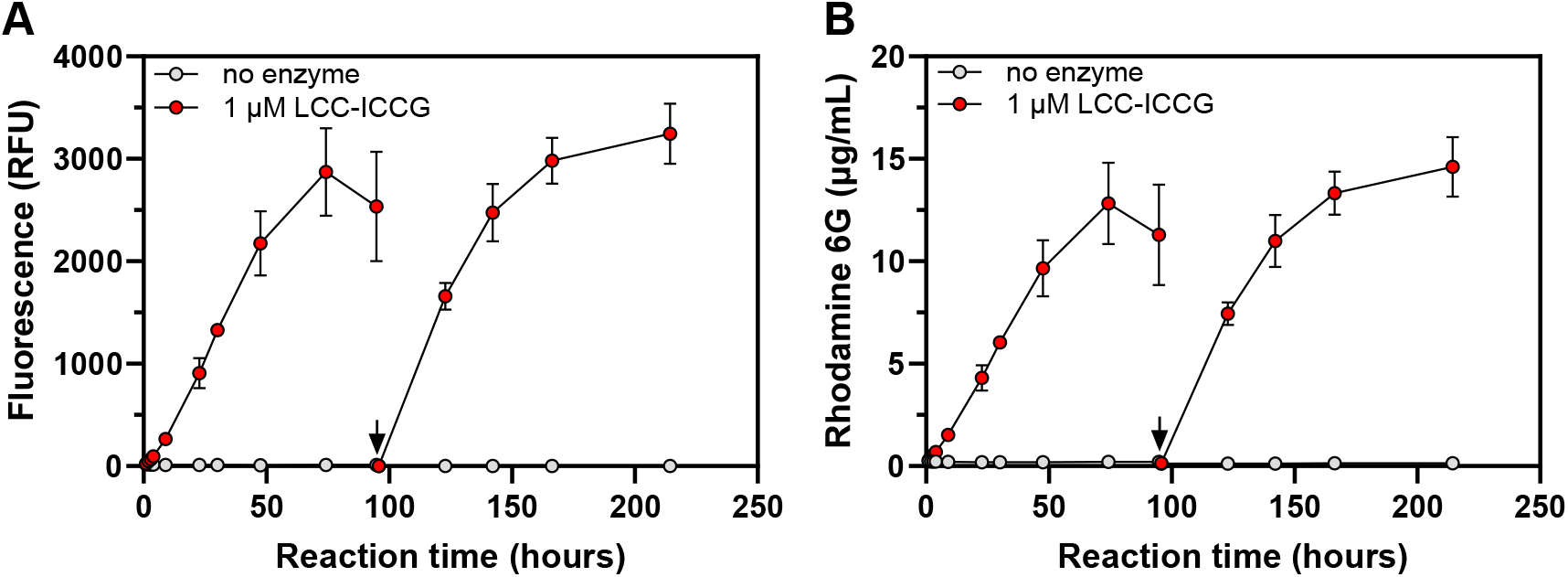
Enzymatic hydrolysis of PET-R6G. **(A)** The fluorescence and **(B)** concentration of free fluorophore in reaction mixtures containing one PET-R6G pellet (∼17 mg; 1.7% w/v solids loading) and 1 µM (28.8 μg) LCC-ICCG (1.7 mg enzyme/g PET). At 95 h, the substrate pellets were transferred to freshly-made reaction mixtures containing buffer and enzyme (indicated by the black arrow). The experiments were conducted in 50 mM Tris-HCl buffer, pH 8.0 at 65 °C. Negative control reactions (“no enzyme”) contained a single PET-R6G pellet and buffer only. Fluorescence was measured at λ_ex/em_ = 530/552 nm after diluting the samples 10-fold with reaction buffer. The concentration of rhodamine 6G was determined according to a standard curve. Error bars indicate standard deviations of triplicate measurements. Source data are provided as a Source Data file.

After 50 hours, the rhodamine 6G release slowed down, indicating either fluorophore depletion or enzyme inactivation. To investigate the latter, the PET-R6G pellets were transferred to freshly prepared reaction mixtures after 95 h of experiment. The resulting progress curves are essentially identical to those in the first reaction (**Fig. 3**), showing that the flattening out of the progress curves is caused by enzyme inactivation rather than rhodamine 6G depletion. Since LCC-ICCG activity is not inhibited by relevant concentrations of free rhodamine 6G (**Fig. S4**), the most likely explanation for reduced activity over time is acidification of the reaction mixture by hydrolysis products. Indeed, an additional control experiment showed that the pH of the reaction mixtures decreased from 8.0 to 4.6 after 96 h of PET-R6G degradation by LCC-ICCG (**Fig. S5**).

The enduring constant rate of fluorophore release shown in **Fig. 3** strongly indicates that the probe is homogeneously distributed within the polymer. In total, ∼26 μg of rhodamine 6G was released into the reaction medium (**Fig. 3B**) after more than 200 hours of incubation. This corresponds to at least 15% PET-R6G solubilization. The actual level of solubilization is higher, since (1) the probe concentration is slightly underestimated due to repeated addition of fresh reaction buffer into the mixtures during sampling (see “Methods” for details), and (2) the actual rhodamine 6G loading in the PET-R6G may be somewhat lower than 1 % (w/w). Most importantly, control reactions lacking the enzyme showed essentially no leakage of rhodamine 6G from PET-R6G, demonstrating the robustness of the assay.

To show that PET-R6G can be used to detect enzymatic activity at lower temperatures (i.e., well below the glass transition point of PET), an additional degradation experiment was carried out at 37 °C, resulting in clear, enzyme-dependent release of the probe, although at a much lower level (∼0.2 µg of rhodamine 6G was released after 19 h at 37°C (**Fig. S6**), compared to ∼4.3 µg after 22 h at 65 °C; **Fig. 3B**).

To confirm that the increase in rhodamine 6G fluorescence observed in the presence of plastic-active enzymes is directly coupled to polymer degradation, the activity of 14 previously reported PETases [14, 16-28] towards PET-R6G was determined in a 24-h endpoint experiment. These enzymes were expressed and purified using a previously described, open-source robotics-enabled pipeline [29]. In this experiment, both the total concentration of soluble aromatic products resulting from PET depolymerization and rhodamine 6G fluorescence were determined (**Fig. 4**). The data revealed correlation between the probe fluorescence and PET depolymerization measured by soluble product release (R^2^ = 0.95). Notably, this experiment was performed at relatively high temperature (70 °C), so it is likely that the observed variation in the performance of PET-active enzymes is partly due to differences in their optimal reaction temperature and stability.

**Figure 4.**
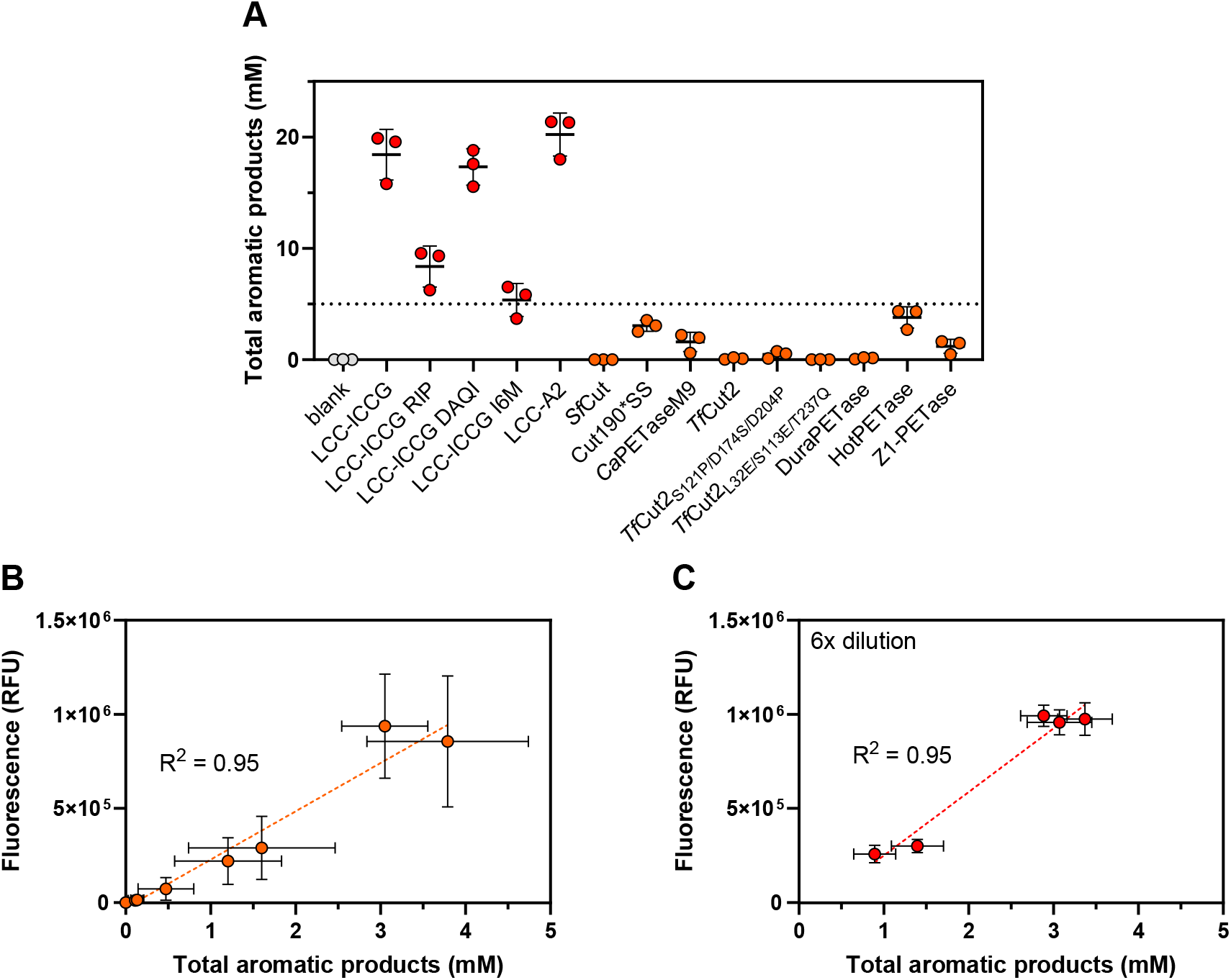
Evaluation of PET-R6G performance using multiple PET-active enzymes. **(A)** The concentration of soluble aromatic products determined by UHPLC after 24 hours of PET degradation and **(B**,**C)** the correlation between the product yield and the rhodamine 6G fluorescence. The experiments were conducted in 0.5 mL 50 mM sodium phosphate buffer, pH 7.5, at 70 °C using a single PET-R6G pellet (∼17 mg, 3.4% w/v solids loading) and 2.5 µg enzyme (0.14 mg/g PET-R6G). Note that the reactions yielding >5 mM soluble aromatic products were diluted 6-fold prior measuring the fluorescence **(C)** to minimize quenching. The product concentrations shown in **(C)** are adjusted to reflect sample dilution. Error bars indicate standard deviations between triplicate measurements. Horizontal lines next to data markers in **(A)** denote mean values. The absolute fluorescence values shown in this figure differ from those in the other experiments, owing to the use of a different fluorometer. Trend lines in **(B)** and **(C)** represent linear fits of the average experimental values performed in GraphPad Prism v. 10.2.0. Source data, including the regression parameters, are provided as a Source Data file.

### Using PET-R6G to assess PETase activity in crude samples

After confirming the consistent performance of PET-R6G in well-defined reactions with purified enzymes, the substrate was validated under conditions relevant for high-throughput screening assays. For this purpose, lysates of *E. coli* BL21 Star (DE3) cells expressing LCC-ICCG were used to degrade PET-R6G, in both a crude form and after removing cell debris by centrifugation (clarified form). Strong fluorescence was observed in reactions containing lysates produced from the LCC-ICCG-expressing *E. coli* strain, whereas almost no fluorescence was detected in control reactions with lysates of *E. coli* devoid of the LCC_ICCG expression plasmid (**Fig. 5A**). Interestingly, the clarified lysates showed lower activity than the crude lysates, suggesting that a fraction of the enzyme co-precipitated with the cell debris. The medium used for cultivating LCC-ICCG-expressing *E. coli* cells also showed substantial activity towards the substrate, which is not surprising given the observed secretion of the enzyme (**Fig. 5B;** note that the enzyme used in our study lacked a signal peptide). Such apparent secretion of recombinant cutinases has been reported before, likely reflecting destabilization of the bacterial cell membrane by the enzyme [30-32]. Our results obtained with the culture medium demonstrate that PET-R6G can potentially be used to set up one-pot experiments for easy activity screening, by growing enzyme-secreting bacteria or fungi, or microbial communities, in the presence of this substrate. The key condition for such experiments to work is the minimal interference of accumulating rhodamine 6G with cell growth. Admittedly, there is evidence for rhodamine 6G toxicity in the literature (see Alford *et al*. for a review [33]), but at the same time, our data obtained by growing BL21 Star (DE3) *E. coli* cells in the presence of this fluorophore show no immediate toxic effect at quite high (100 µg/mL) probe concentration (**Fig. 6**). Such concentration is expected to be reached upon more than 50% solubilization of a single PET-RG pellet in 1 mL reaction.

**Figure 5.**
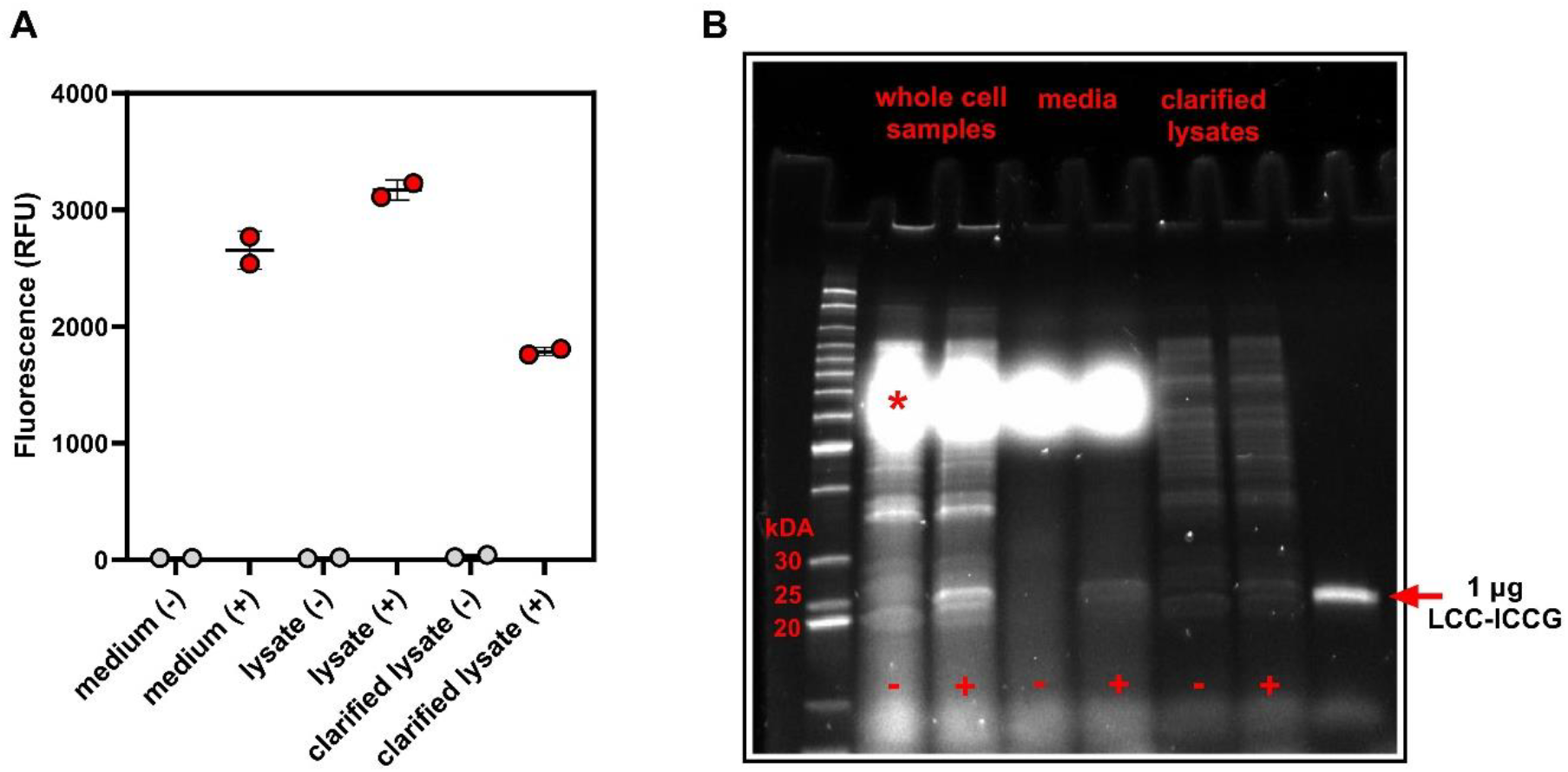
Detecting PETase activity in crude biological samples. **(A)** Degradation of PET-R6G observed in experiments with *E. coli* lysates and cell-free culture media and **(B)** results of SDS-PAGE analysis of these preparations compared to whole cell samples (i.e., total culture prior to lysis). LCC-ICCG-containing samples are marked with “+”, whereas negative control samples are labeled with “-”. The dilution factor was the same across all experimental conditions, meaning that the band intensities reflect the true differences in protein abundance. The image color was inversed to improve visual clarity. The bright high molecular mass bands (annotated with “*”) originate from medium compounds interfering with the stain-free protein detection system. The PET-R6G degradation experiments were carried out in duplicates for 60 hours at 60 °C, 500 RPM, in 0.5 mL of 50 mM Tris-HCl, pH 8.0 (1 pellet per reaction). Error bars indicate standard deviations; horizontal lines denote average values. Source data are provided as a Source Data file.

**Figure 6.**
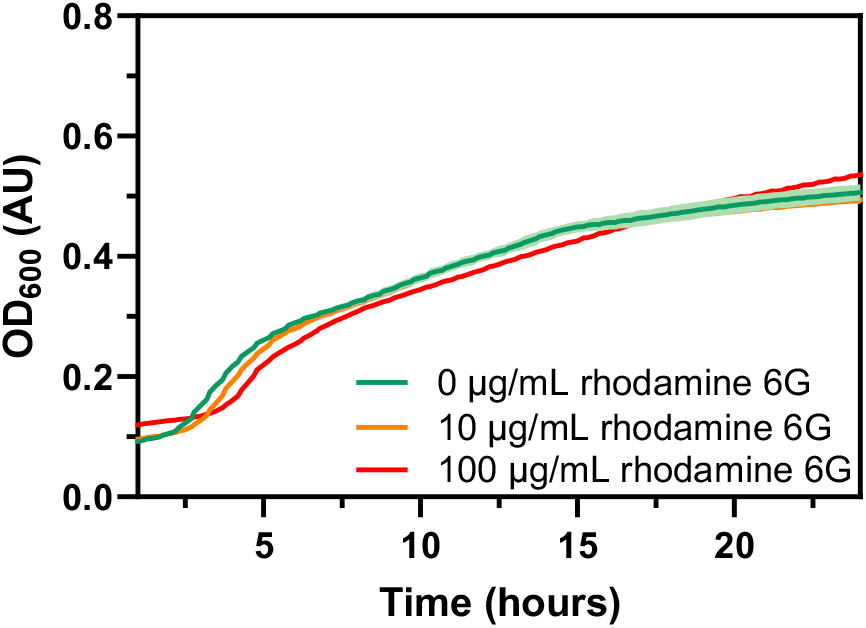
Effect of rhodamine 6G on growth of *E. coli*. Growth curves of BL21 Star (DE3) cells incubated in LB medium at 37 °C in the presence of 0, 10, or 100 µg/mL rhodamine 6G. Error envelopes (visible for the green trace only) indicate standard deviations between triplicate measurements. The experiment was conducted in a microtiter plate with a <1 cm optical path. Source data are provided as a Source Data file.

### Preparation and evaluation of polyethylene- and nylon-based fluorophore-doped materials

Encouraged by the results obtained with PET-R6G, we produced an additional selection of blended plastics with high density polyethylene (HDPE), nylon 6, and nylon 6,6 using a similar compounding/extrusion procedure as before (see **Fig. S1** for the extrusion temperatures and **Table S1** for the DSC analysis of the source and resulting materials). Unsurprisingly, the performance of the new materials in passive diffusion tests was dependent on the polymer type (**Fig. 7**).

**Figure 7.**
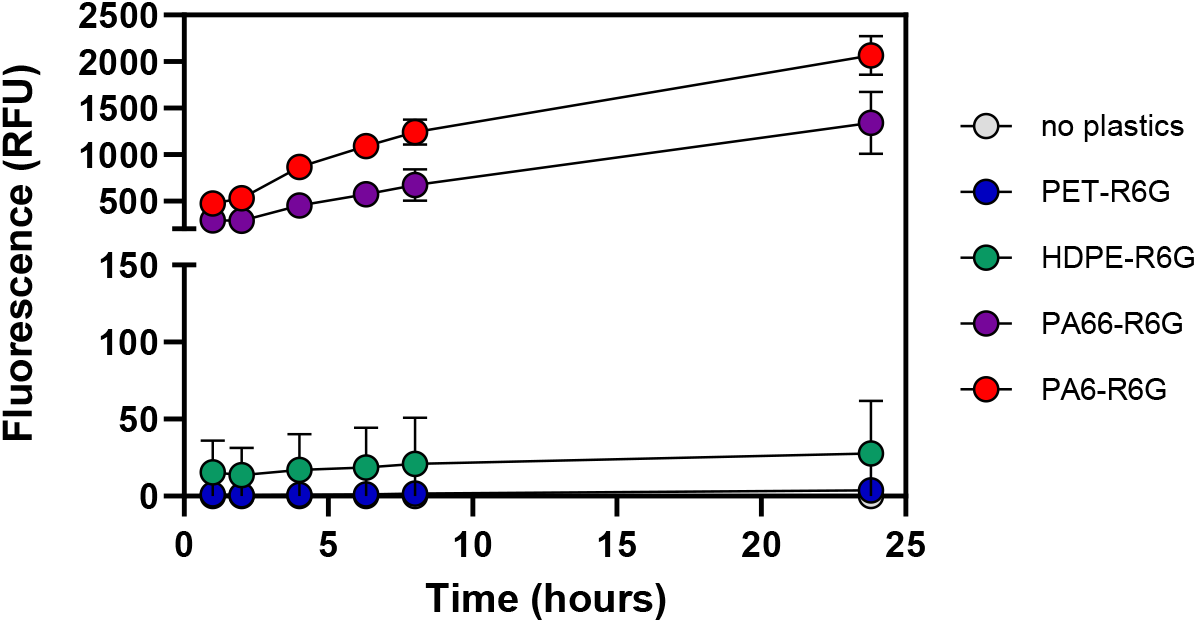
Leaching of rhodamine 6G from various blended plastics. The curves show the increase in fluorescence observed during the incubation of blended materials in 1 mL of 50 mM Tris-HCl, pH 8.0, at 65 °C, 1000 RPM (1 pellet per experiment). Error bars indicate standard deviations between triplicate measurements. Note that the fluorescence values (λ_ex/em_ = 530/552 nm) were obtained after 10-fold dilution with buffer. All rhodamine containing materials were prepared at 1% (w/w) rhodamine 6G loading, but the actual loading of the resulting materials may have been lower. Source data are provided as a Source Data file.

Rhodamine 6G-loaded HDPE pellets (HDPE-6G) displayed a low degree of fluorophore leaching, similar to the PET-R6G. Conversely, both nylon 6 and nylon 6,6-based blends (PA6-R6G and PA66-R6G, respectively) showed rapid release of rhodamine 6G into the reaction buffer, with continuous leaching from the nylon substrates during the experiment (∼24 hours).

To date, there is no compelling evidence of enzymes capable of degrading non-pretreated polyolefins (such as polyethylene) that can be used to set up positive control experiments with HDPE-R6G. Yet, it is tempting to suggest that the low background signal observed with this material could allow using it to screen for such activity in future. Conversely, several families of nylonases are known and sufficiently characterized [34], which permitted testing the blended nylon substrates. Of note, the achievable conversion extents for the best nylonases reported to date are still low, ∼1 wt %, and amorphous film substrates are commonly used to study nylon depolymerization rather than pelletized plastics [34, 35]. In our experiments, a top-performing engineered thermostable nylonase from *Kocuria sp*. (NylC_K_-TS; an N-terminal nucleophile (Ntn) hydrolase) [34] capable of acting on nylon 6 was used as a model enzyme. After more than 10 days of incubation with 1 µM nylonase, no enzyme-dependent release of fluorophore from PA6-R6G was observed (**Fig. 8**), likely due to the combination of a high background signal and low conversion of this crystalline PA6-R6G substrate (37.3 ± 0.5%; **Table S1**).

**Figure 8.**
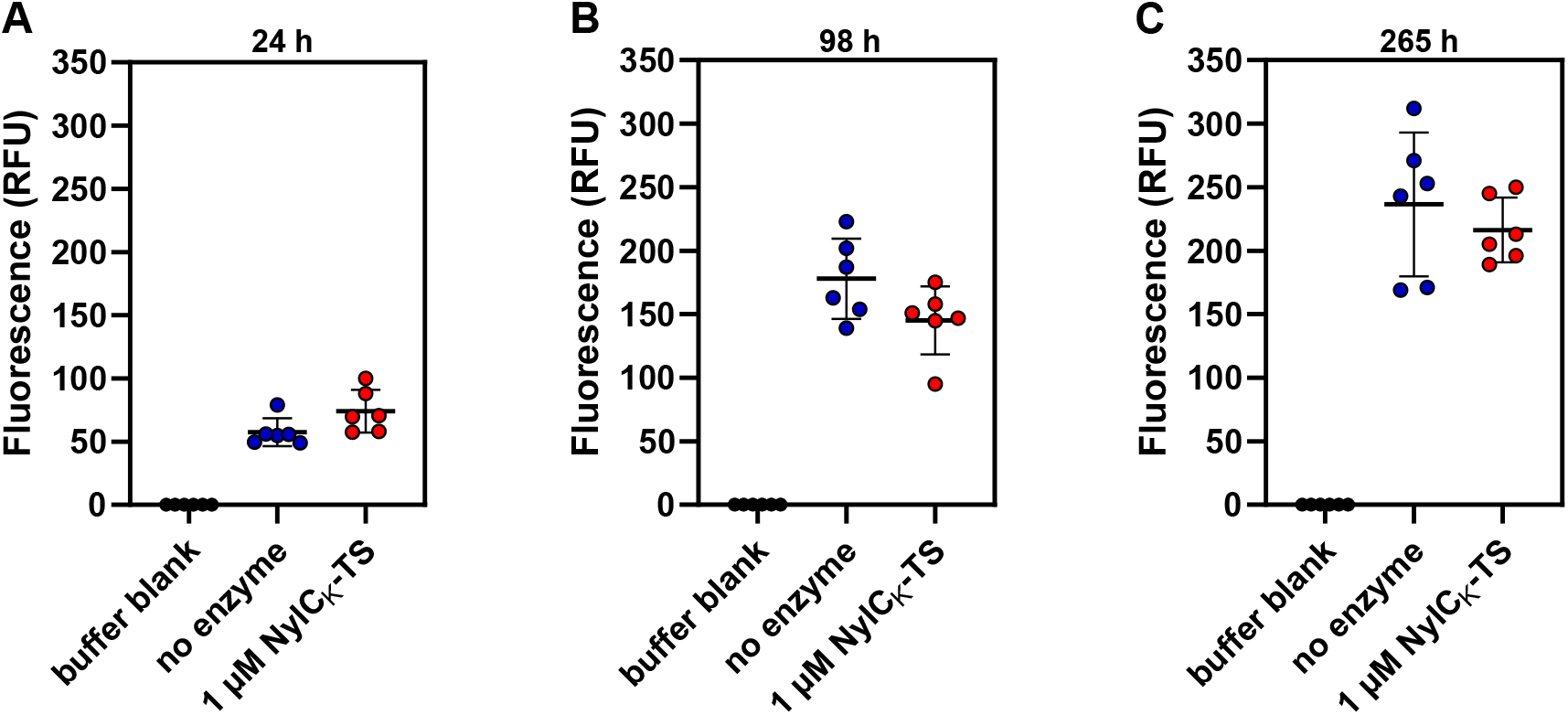
Rhodamine 6G fluorescence in reactions with PA6-R6G in the presence or absence of NylC_K_-TS. From left to right, the three panels show the observed fluorescence (λ_ex/em_ = 530/552 nm) in 1 ml reaction mixtures containing PA6-R6G (∼11 mg; 1.1 % w/v solids loading) after 24, 98 and 265 h incubation in the presence or absence of 1 µM (39 µg; 3.5 mg enzyme/g PA6-R6G) nylonase. The experiments were carried out at 65°C, 500 RPM in 50 mM sodium phosphate buffer, pH 7.4, supplied with 150 mM NaCl. Error bars indicate standard deviations between replicates (*n* = 6). Horizontal lines denote average values. Note that fluorescence was measured after diluting the samples 100-fold with 50 mM Tris-HCl, pH 8.0. **Fig. S7** shows the results of product analysis by MALDI-ToF MS performed with non-diluted endpoint samples. The buffer blank signals shown in all three panels represent the same experiment (*n* = 6). Source data are provided as a Source Data file.

To further investigate the potential effects of NylC_K_-TS on PA6-R6G, the soluble fractions of the resulting reaction mixtures were analyzed using matrix-assisted laser desorption/ionization (MALDI) mass spectrometry (**Fig. S7**). The mass spectrometry data indicated enzyme-dependent formation of linear nylon-6 oligomers with various degrees of polymerization (DP), confirming nylonase activity. However, substantial amounts of cyclic oligomers were detected in both experimental and control samples (DP2-DP6), complicating interpretation of the data. Cyclic oligomers are well-known contaminants in nylon 6 [36], representing side-products of polymerization. Importantly, the multi-step washing performed as a part of substrate preparation (see “Methods” for more details) did not remove this contamination, suggesting that cyclic oligomers were continuously leaching from PA6-R6G during the experiment. NylC_K_-TS is known to act on such cyclic substrates converting them to linear oligomers [34], which is also suggested by the mass spectra (**Fig. S7**), indicating some depletion of the cyclic pentamer and hexamer upon enzyme addition. All in all, we cannot exclude that some of the observed linear reaction products originate from the hydrolysis of low molecular weight cyclic contaminants.

## Discussion

The production of fluorophore-loaded blends to track enzymatic deconstruction of plastics is simple and appealing. If a uniform distribution of a water-soluble probe can be achieved within a polymer matrix, the resulting system can be, in principle, used to obtain quantitative information on polymer degradation, regardless of the chemical properties of all potential (soluble, insoluble, or volatile) cleavage products. Despite the potential advantages, this approach is not common due to time-consuming substrate preparation protocols.

This study presents a convenient and flexible method for producing a variety of rhodamine 6G-loaded substrates on at least a 200-gram scale (∼12,000 substrate pellets). Melt blending followed by extrusion is a well-controlled compounding process that can be easily reproduced and scaled up when needed. Given the relatively low price of rhodamine 6G, very large batches of blended plastics can be prepared and stored, allowing large high-throughput screens with minimal substrate production efforts. Pelletized substrates are easy to handle compared to setting up reactions with films or powders, especially given the complications associated with the accumulation of electrostatic charge on the latter.

Our results clearly show that PET-R6G can be used to assess enzymatic activity in a high-throughput fashion. In time-course experiments, the release of the fluorophore was linear over time for many hours suggesting uniform distribution of the probe within the PET matrix, which is further confirmed by the correlation between the fluorescence and UHPLC data.

Despite the popularity of rhodamine 6G as a fluorescent probe, little information is available on its physical properties such as melting temperature. While some literature sources claim the melting temperature to be as high as 290 °C [37], product data from vendors suggest that is could be lower (e.g., 230 °C; Carl Roth, Karlsruhe, Germany, article number 0749.2). The compounding process applied to produce PET-R6G involved a temperature of 250 °C, so it is possible that the consistent performance of our substrate is due to melting of rhodamine 6G, allowing for a truly homogeneous PET/probe mixture during extrusion. In retrospect, it is clear that rhodamine 6G is sufficiently stable during the melt blending, so the 1% (w/w) fluorophore load is excessive. Lower loads (e.g., 0.1%) could likely be used to minimize fluorescence quenching (i.e., the need for diluting samples), without affecting the method sensitivity. Importantly, our compounding process had a limited effect on the crystallinity of all the resulting materials (**Table S1**), except for PET, which exhibited a significant decrease in the crystallinity after melt blending (4.8 ± 0.9% compared to 33.3 ± 0.7% in the source material). Producing PET-R6G with a higher degree of crystallinity is of interest, and could be achieved by optimizing the extrusion conditions.

It is worth noting that the aromaticity of PET degradation products makes these compounds easy to detect and quantify by simply measuring the increase in UV absorbance around 240 nm. Yet, using PET-R6G offers an important advantage. Following rhodamine 6G fluorescence (λ_ex/em_ = 530/552 nm) allows for remarkably low background signals in crude biological samples, such as bacterial lysates or culture media, as shown in our study. Robust signal-to-noise ratios and the high specificity of the signal (compared to UV-absorbance) open up opportunities for even faster/simpler experiments, such as cultivating PETase-secreting bacteria or fungi, or even microbial consortia, in the presence of PET-R6G.

In contrast to PET-R6G, the performance of PA6-R6G and PA66-R6G substrates was not satisfactory. Rapid and continuous diffusion of rhodamine 6G from these polyamides resulted in high background signals. This, combined with the low activity of NylC_K_-TS on crystalline (37.3 ± 0.5%) PA6-R6G under our experimental conditions, likely explains why no enzyme-dependent release of the fluorophore was observed. One of the simplest solutions to the high background problem faced with nylon substrates could be using a bulkier rhodamine 6G derivative to slow down diffusion. Multiple candidates are available for consideration (see Beija et al. for a review [9]); however, any potential modification to rhodamine 6G will not only affect the fluorescence and solubility of the probe, but also its price. Given this, the probably most sensible strategy for impeding passive diffusion could be using commercially available fluorescent carbon nanodots as tracer species. Such an approach has been successfully applied in the production of PCL films [8].

As for our HDPE-based blended material, a negligibly low background signal was observed during the passive diffusion tests. Ironically, there is no enzyme that can be used as a positive control to further validate the performance of this substrate. Despite several claims that some redox enzymes can break down non-pretreated non-hydrolyzable plastics such as PE, poly(vinyl chloride) or polypropylene, this idea remains controversial [1, 38]. There are indications that many, if not all, of previously published claims regarding enzymes with the ability to depolymerize non-hydrolyzable plastics, may have been based on misinterpreted results [39]. Recent attempts at replicating some of the most prominent studies with such claims were not successful [40, 41]. Conversely, there is a growing amount of literature showing that ester bonds can be incorporated into the polyethylene backbone via photo, chemical, or chemo-enzymatic pre-treatment [42-44], allowing for subsequent hydrolysis by esterases. It could be of interest to produce fluorophore-loaded variants of such pre-oxidized PE materials to allow rapid screening for degradative enzymatic activities and reaction conditions.

## Methods

### Chemicals and materials

Chemicals were sourced from Sigma-Aldrich (St. Louis, MO, USA) unless specified otherwise. Low-melting point PET (CumaPET L04 100) was obtained from DuFor Resins BV (Zevenaar, the Netherlands). Nylon-6 and Nylon-6/6 resins (Technyl® Shape C 402M NC; Technyl® Safe A 402FC NC, respectively) were sourced from DOMO Chemicals (Gent, Belgium). High density polyethylene (grade SHA7260) was sourced from Braskem Europe (Wesseling, Germany). A stock solution of 4-Nitrophenyl acetate was prepared at 100 mM concentration in EtOH and stored at -20 °C. Rhodamine 6G chloride powder was kept at room temperature protected from direct light to avoid photobleaching.

### Preparation of fluorophore-loaded plastics

PET-R6G, HPDE-R6G, PA6-R6G and PA66-R6G pellets were produced by melt blending of milled plastic granules with rhodamine 6G using a Euro Lab Prism 16 twin-screw extruder (Thermo Fisher Scientific, Waltham, MA, USA) with an L/D ratio of 25:1. The appropriate amounts of polymer and rhodamine 6G powders were mixed to achieve 1% w/w fluorophore loading before processing with an extruder at 500 RPM. The extrusion temperature varied for different resins. PE, PET, PA6, and PA66 were processed at 190°C, 250°C, 230°C, and 265°C, respectively. The extrudate strands were either air-cooled (PA6-R6G, PA66-R6G) or water-cooled (PE-R6G, PET-R6G) and then pelletized with an SGS 50-E pelletizer (C.F. Scheer & Cie, Stuttgart, Germany). The resulting pellets were stored at room temperature away from light. In total, 200 g of each substrate was prepared.

Prior to the experiments, additional washing was applied to remove fluorophore adsorbed on the surface of blended materials. The washing was performed by vortexing 4 g of pellets in 40 mL of milli-Q water or 100% EtOH for 5 minutes and discarding the liquid fractions. The number of wash steps and washing liquid depended on the blend type: the PET-R6G pellets were washed three times in Milli-Q water followed by one wash in 100% EtOH, the HDPE-R6G pellets were washed three times in 100% EtOH and the PA6-R6G and PA66-R6G pellets were washed five times in Milli-Q water, followed by two washes in 100% EtOH. The pellets were air-dried after the washing and stored at room temperature away from light until further use.

### Determination of crystallinity in polymers

The crystallinity in virgin and blended materials was assessed by differential scanning calorimetry using a DSC250 calorimeter (TA instruments, New Castle, DE, USA). Plastic samples were encapsulated into sealed aluminium containers and subjected to heating according to the following protocol:

PET/PET-R6G: from 20 °C to 280 °C at a rate of 10°C/min followed by an isothermal hold of 5 min. HDPE/HDPE-R6G: from 20 °C to 200 °C at a rate of 10°C/min followed by an isothermal hold of 5 min. PA6/PA6-R6G: from 20 °C to 260 °C at a rate of 10°C/min followed by an isothermal hold of 5 min. PA66/PA66-R6G: from 20 °C to 300 °C at a rate of 10°C/min followed by an isothermal hold of 5 min. The percentage of crystallinity was calculated using the following equation:

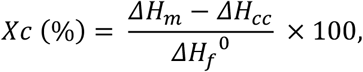

where *ΔH*_*m*_ is the observed enthalpy of melting, *ΔH*_*cc*_ is the observed enthalpy of cold crystallization and *ΔH*_*f*_^*0*^ is a reference value for the enthalpy of melting of 100% crystalline material (*ΔH*_*f*_^*0*^ values of 140 J g^-1^, 290 J g^-1^, 190 J g^-1^ and 290 J g^-1^ were used for PET, HDPE, nylon 6 and nylon 6,6, respectively). The analysis was carried out in duplicates. Note that the cold crystallization term *ΔH*_*cc*_ was omitted when calculating the crystallinity of all plastics except for PET, as cold crystallization was observed for this polymer only.

### Protein expression and purification

Previously obtained plasmids encoding for LCC-ICCG [14] and NylC_K_-TS [34] with C-terminal His-tags (pET-21(b)+ and pET-28(a)+, respectively) were used for protein expression in *E. coli*. The detailed information on these vectors has been previously deposited at the AddGene repository [https://www.addgene.org/Gregg_Beckham/].

Chemically competent BL21 Star (DE3) *E. coli* cells (Thermo Fisher Scientific; Waltham, MA, USA) were transformed with the corresponding plasmids according to the supplier’s protocol. Single colonies were picked from LB agar plates containing 50 µg/mL kanamycin (pET-28(a)+) or 100 µg/mL ampicillin (pET-21(b)+) after overnight incubation at 37 °C and were further cultivated in LB medium overnight at the same temperature (200 RPM). These starter cultures were used to inoculate 0.5 L of LB medium supplied with appropriate antibiotics. The cells were then incubated at 37 °C, with shaking at 200 RPM, until an OD_600_ of approximately 0.6 was reached. Next, the cultures were cooled down to 25 °C and protein expression was induced by adding isopropyl thiogalactopyranoside (IPTG) to 0.5 mM final concentration. The cells were harvested by centrifugation (6,000 x g for 15 min) after 24 hours of growth and resuspended in 30 mL of binding buffer (50 mM Tris-HCl, pH 8.0, supplied with 5 mM imidazole and 500 mM NaCl). The cells in the resulting suspensions were lysed by sonication using a VibraCell ultrasonic disintegrator equipped with a micro tip probe (Sonics, Newtown, CT, USA) for 10 min in 5 s steps followed by 5 s pauses at 29% amplitude. The lysates were clarified by centrifugation at 20,000 x g for 15 min at 4 °C and filtered through a 0.22 µm syringe filter. The enzymes were purified from lysates by immobilized metal affinity chromatography using a 5 mL Ni-charged HisTrap FF column (Cytiva, Marlborough, MA, USA). The proteins were eluted with a linear gradient of imidazole (5 − 500 mM) in binding buffer (20 column volumes at 2.5 mL/min flow rate). Protein preparations were analyzed with SDS-PAGE. Imidazole and NaCl were removed from enzyme stock solutions by multiple rounds of concentration-dilution using 50 mM Tris-HCl buffer (pH 8.0) and Vivaspin ultrafiltration tubes (10 kDa MWCO; Sartorius, Göttingen, Germany). Protein concentrations were determined by UV spectroscopy (λ = 280 nm) using theoretical extinction coefficients calculated with ProtParam tool [45].

### Following enzymatic degradation of PET-R6G by LCC-ICCG

To study the effect of LCC-ICCG on PET-R6G, pre-washed substrate pellets (1 pellet per reaction; ∼17 mg PET-R6G) were incubated in 1 mL of 50 mM Tris-HCl, pH 8.0, containing or lacking 1 µM enzyme, at 65 °C, 1000 RPM, using a Thermomixer (Eppendorf, Hamburg, Germany). To determine the release of rhodamine 6G into the reaction mixture, 20 µL aliquots were taken at various time points, diluted to 200 µL with the reaction buffer and loaded into a non-transparent 96-well microtiter plate. The fluorescence was measured using a VarioSkan Lux plate reader (Thermo Fisher Scientific; Waltham, MA, USA) at the excitation/emission wavelength of 530/552 nm (1 nm bandwidth; 100 ms integration time). To compensate for the volume loss, 20 µL of reaction buffer containing or lacking 1 µM LCC-ICCG was added into the reaction mixtures immediately after each sampling. After approximately 95 hours of incubation, the substrate pellets were transferred into 1 mL of freshly prepared reaction buffer, containing or lacking 1 µM enzyme, to restart the experiment. The reactions were carried out in triplicates. The concentration of released rhodamine 6G was calculated according to a standard curve by performing non-linear fit of the calibration and experimental data using the four-parameter logistic (4PL) model. Data processing was performed in GraphPad Prism v. 10.2.0. An additional set of enzymatic and control reactions (*n* = 5) was set up at lower temperature and slower agitation (37 °C, 500 RPM). 200 µL aliquots were taken from the reaction mixtures after 19 hours and fluorescence was measured in these undiluted samples.

### Evaluation of PET-R6G performance using a collection of PET-active enzymes

LCC-ICCG (prepared separately for this experiment; see below), LCC-ICCG RIP, LCC-ICCG DAQI, LCC-ICCG I6M, LCC-A2, *Sf*Cut, Cut190*SS, *Ca*PETaseM9, *Tf*Cut2, *Tf*Cut2_S121P/D174S/D204P_, *Tf*Cut2_L32E/S113E/T237Q_, DuraPETase, HotPETase, and Z1-PETase were expressed and purified using a previously described robotic platform [29]. Briefly, plasmids containing the codon optimized sequences cloned into pCDB179 (gifted to Addgene by Christopher Bahl, #91960) were transformed into chemically competent C41(DE3) *E. coli*. Protein expression occurred in 2 mL autoinduction cultures supplemented with 100 µg/mL kanamycin and 1X trace metals (Teknova, Hollister, CA, USA) and incubated with shaking for 2 h at 37 °C followed by 40 h at 18 °C. Cells were harvested by centrifugation and then lysed by resuspending in 20 mM Tris-HCl, pH 8.0, 300 mM NaCl, 5 mM imidazole, 1% n-octyl β-D-glucopyranoside supplemented with 0.1 mg/mL DNase I and 1 mg/mL lysozyme, followed by shaking for 1 h. Ni-charged magnetic beads (GenScript; Piscataway, NJ, USA; catalog number L00295) were introduced to bind the His-tagged recombinant enzymes and then washed with 20 mM Tris-HCl, pH 8.0, 300 mM NaCl to remove cell debris. His-tagged Cth SUMO protease was used to remove the target proteins from the magnetic beads by cleaving the SUMO/His-tags, and the concentrations of purified enzymes were determined using the Pierce Rapid Gold BCA Protein Assay Kit (Thermo Fisher Scientific; Waltham, MA, USA).

For activity measurements, 2 mL deep-well 96-well plates were loaded with one pellet of PET-R6G and 475 µL of buffer (50 mM sodium phosphate buffer, pH 7.5), sealed with aluminum sealing tape, and incubated at 70 °C for 2 hours to preheat the assays plates. 25 µL of enzyme solution at 0.1 mg/mL (or 25 µL of buffer solution lacking the enzyme) was added to start the experiments. The plates were sealed with a heat sealer and incubated at 70 °C with shaking at 250 rpm (19 mm orbit). For fluorescence measurements, 100 µL aliquots were transferred to flat bottom 96-well plates for analysis in a BioTek Synergy H1 plate reader (Agilent Technologies, Santa Clara, CA, USA) using an excitation/emission wavelength of 530/552 nm. To minimize quenching at high fluorophore concentrations, high yielding samples (> 5 mM total soluble aromatic products according to UHPLC analysis) were diluted six-fold prior to measuring the fluorescence.

### UHPLC detection and quantification of PET degradation products

Quantification of terephthalic acid, mono-(2-hydroxyethyl)terephthalic acid and bis-(2-hydroxyethyl)terephthalic acid) was performed using ultra-high performance liquid chromatography (UHPLC) as described in a prior study [20]. In short, both standards and samples were analyzed with an Infinity II 1290 ultra-high-performance liquid chromatography system (Agilent Technologies; Santa Clara, CA, USA) equipped with a G7117A diode array detector. The products were separated using a Zorbax Eclipse Plus C18 Rapid Resolution HD column (Agilent Technologies) and a gradient of methanol in phosphoric acid. Detection was carried out at 240 nm, and calibration curves were generated to quantify each target compound. Samples above the detection range were diluted six-fold and final reported concentration values corrected for dilution.

### Inhibition of LCC-ICCG by rhodamine 6G

LCC-ICCG inhibition by rhodamine 6G was assessed using 4-nitrophenyl acetate as a model substrate. Reaction mixtures (1 mM 4-nitrophenyl acetate in 50 mM Tris-HCl, pH 8.0) containing 0, 10, 100 or 1000 µg/mL fluorophore were loaded into a 96-well microtiter plate (100 µL per well). The experiments were initiated by adding enzyme to 0.1 µM final concentration or by adding the same volume of milli-Q water. The hydrolysis of 4-nitrophenyl acetate was followed at the room temperature by measuring the absorbance of 4-nitrophenol at 400 nm using a VarioSkan Lux plate reader (Thermo Fisher Scientific; Waltham, MA, USA). The reaction rates (mAU * min^-1^) were determined using linear parts of the resulting progress curves.

### Detection of PETase activity in *E. coli* lysates and culture medium

Chemically competent BL21 Star (DE3) *E. coli* cells (Thermo Fisher Scientific; Waltham, MA, USA) were transformed with a pET-21(b)+ vector encoding for LCC-ICCG, spread on an LB agar plate containing 100 µg/mL ampicillin and incubated overnight at 37 °C. A single colony was transferred into 20 mL of fresh LB medium supplied with the same antibiotic at the same concentration. The cells were incubated overnight at 37 °C, 200 RPM. To generate a control culture, untransformed BL21 Star (DE3) *E. coli* cells were grown overnight in 20 mL of LB medium (37 °C, 200 RPM) in the absence of antibiotic. Next, both cultures were induced by adding IPTG to 1 mM final concentration and further incubated for 3 more hours at 37 °C and 200 RPM. Cells were harvested by centrifugation at 4,000 x g for 15 minutes, resuspended in 20 mL of 50 mM Tris-HCl, pH 8.0, and subjected to ultrasonic lysis as described above (see “Protein expression and purification”). Half of the volume of each lysate was clarified by centrifuging at 20,000 x g for 15 minutes, whereas the other half remained unprocessed. 50 µL of crude lysates, clarified lysates and culture medium were diluted 10-fold with 50 mM Tris-HCl, pH 8.0, and co-incubated with PET-R6G (1 pellet per reaction) at 60°C, 500 RPM for 64 hours before measuring fluorescence.

### Rhodamine 6G toxicity assay

BL21 Star (DE3) *E. coli* cells were cultivated overnight at 37 °C, 200 RPM in 20 mL LB medium and diluted to OD_600_ = 0.01 using a fresh batch of the same medium. 100 µL aliquots were transferred from the diluted culture to a 96-well microtiter plate (1 aliquot per well). To each well, 80 µL of fresh LB medium was added followed by 20 µL of aqueous rhodamine 6G solutions (or Milli-Q water) to yield a final fluorophore concentration of 0, 10 or 100 µg/mL (3 replicates for each fluorophore concentration). The plate was sealed with a transparent film, which was punctured with a syringe needle on top of each well once to allow for gas exchange. The cell growth experiment was carried out at 37°C for 33 h in a Varioskan Lux plate reader (Thermo Fisher Scientific; Waltham, MA, USA), with shaking at 180 RPM for 10 seconds every 5 minutes and OD_600_ recording every 10 min.

### Passive diffusion of rhodamine 6G from various plastics

For each type of blended polymer material, 1 pellet was transferred into a 1.5 mL microcentrifuge tube containing 1 mL of 50 mM Tris-HCl, pH 8.0, and incubated for 23 hours at 65 °C, 1000 RPM. 20 µL aliquots were taken at different time points and diluted 10-fold with the same buffer in a 96-well non-transparent microtiter plate prior to measuring the fluorescence using a Varioskan Lux plate reader. The experiments were carried out in triplicates.

### Probing the degradation of PA6-R6G by NylC_K_-TS

To assess the capacity of PA6-R6G to release rhodamine 6G upon enzymatic degradation, the substrate pellets (1 pellet per 500 µL reaction) were incubated in 50 mM sodium phosphate buffer, pH 7.4, supplied with 150 mM NaCl and containing (or lacking) 1 µM NylC_K_-TS. The experiments were carried out at 65°C, 500 RPM for 265 hours. 2 µL aliquots were taken at various time points and diluted 100-fold with 50 mM Tris-HCl, pH 8.0, in a 96-microtiter plate before measuring fluorescence using a Varioskan Lux plate reader. The reaction mixtures at the 265-h endpoint were analyzed by MALDI-ToF MS by mixing 1 µL sample with 1 µL of matrix solution (9 mg/mL 2,5-dihydroxybenzoic acid in 30% (v/v) acetonitrile) on the surface of an MTP 384 ground steel target plate (Bruker Daltonics GmbH, Bremen, Germany). The target plate was air-dried and the sample was analyzed with a UltrafleXtreme mass spectrometer (Bruker Daltonics) operated in positive mode (200-750 m/z range) and controlled by flexControl 3.4 software.

## Acknowledgements

The work of AAS, ELT, RRC, VGHE, and GVK was supported by the Research Council of Norway under Grant no. 326975 (Enzyclic), with additional support from the SmartPlast sustainability arena of the Norwegian University of Life Sciences. This work was authored in part by the National Renewable Energy Laboratory for the U.S. Department of Energy (DOE) under Contract No. DE-AC36-08GO28308. Funding was provided in part to BNB and GTB by the U.S. Department of Energy, Office of Energy Efficiency and Renewable Energy, Advanced Materials and Manufacturing Technologies Office (AMMTO) and Bioenergy Technologies Office (BETO). This work was performed as part of the Bio-Optimized Technologies to keep Thermoplastics out of Landfills and the Environment (BOTTLE) Consortium and was supported by AMMTO and BETO under Contract DE-AC36-08GO28308 with the National Renewable Energy Laboratory (NREL), operated by Alliance for Sustainable Energy, LLC. This material is also based upon work supported by the U.S. Department of Energy, Office of Science, Office of Biological and Environmental Research (BER), Genomic Science Program under Award Number DE-SC0022024 to BNB and GTB. The views expressed in the article do not necessarily represent the views of the DOE or the U.S. Government. The U.S. Government retains and the publisher, by accepting the article for publication, acknowledges that the U.S. Government retains a nonexclusive, paid-up, irrevocable, worldwide license to publish or reproduce the published form of this work, or allow others to do so, for U.S. Government purposes.

## Contributions

AAS designed and performed experiments, analyzed the data, wrote the first draft, and contributed to manuscript editing. BNB and ELT designed and performed experiments, analyzed the data, contributed to data interpretation, and manuscript editing. RRC designed experiments, contributed to data interpretation and manuscript editing. GTB and VGHE conceptualized the work, contributed to data interpretation, and manuscript editing. GVK supervised the project, conceptualized the work, contributed to data interpretation, and manuscript editing.

