## Supplementary information for "Production and evaluation of fluorophore-doped polymer substrates to screen for plastic-degrading enzymes"

**Table S1. Melting temperatures and crystallinity of the materials used in this study.** The data were acquired in duplicates using differential scanning calorimetry (DSC).

| Material | Melting temperature, °C | Degree of crystallinity, % |
|---|---|---|
| PET | 219.7 ± 2.1 | 33.3 ± 0.7 |
| PET-R6G | 231.9 ± 0 | 4.8 ± 0.9 |
| HDPE | 131.4 ± 0.4 | 63.6 ± 0.2 |
| HDPE-R6G | 132.8 ± 0.6 | 61.2 ± 0.4 |
| PA6 | 216.6 ± 0.9 | 35.4 ± 0.5 |
| PA6-R6G | 215.9 ± 0.2 | 37.3 ± 0.5 |
| PA66 | 260.4 ± 0.1 | 33.7 ± 0.7 |
| PA66-R6G | 261.4 ± 0.5 | 32.3 ± 0.1 |

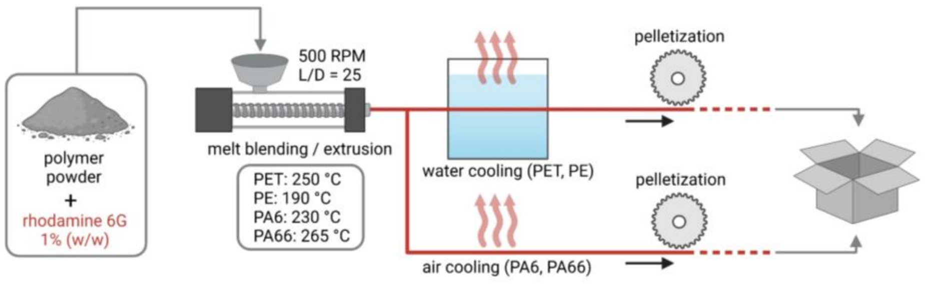


**Figure S1. Schematic representation of the melt blending process used to create fluorophore-doped materials in this study.** The figure was created with BioRender.

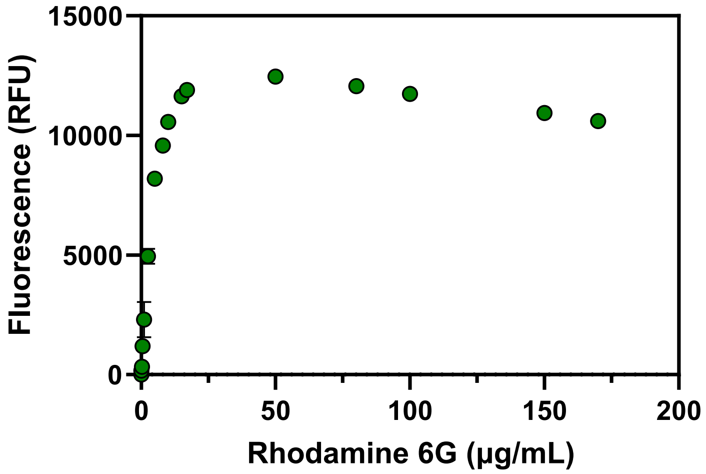


**Figure S2. Fluorescence of standard solutions of rhodamine 6G.** The fluorescence data was obtained using solutions with various concentrations of free rhodamine 6G ($\lambda_{ex/em}$ = 530/552 nm) in 50 mM Tris-HCl buffer, pH 8.0. Error bars indicate standard deviations between triplicate measurements and are in most cases hidden behind the data markers. Note that increasing the fluorophore concentration beyond 50 µg/mL results in a decrease in fluorescence. Source data are provided as a Source Data file.

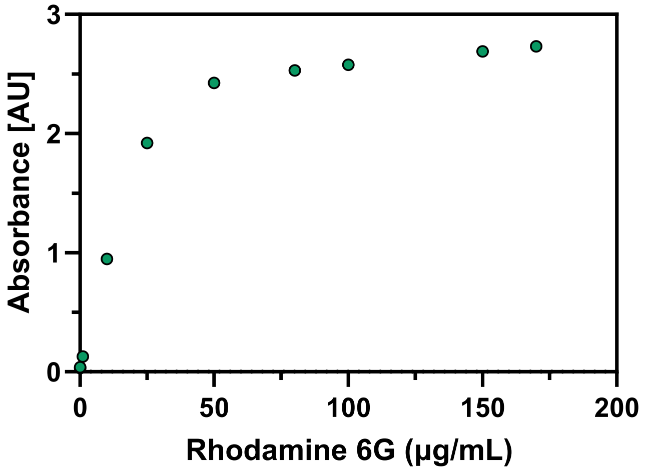


**Figure S3. Light absorbance of rhodamine 6G standard solutions.** The data was acquired at 530 nm in a 96-well microtiter plate with a <1 cm optical path. 200 µL samples were prepared in triplicates in 50 mM Tris-HCl, pH 8.0. Error bars indicating standard deviations are hidden behind the data markers. Source data are provided as a Source Data file.

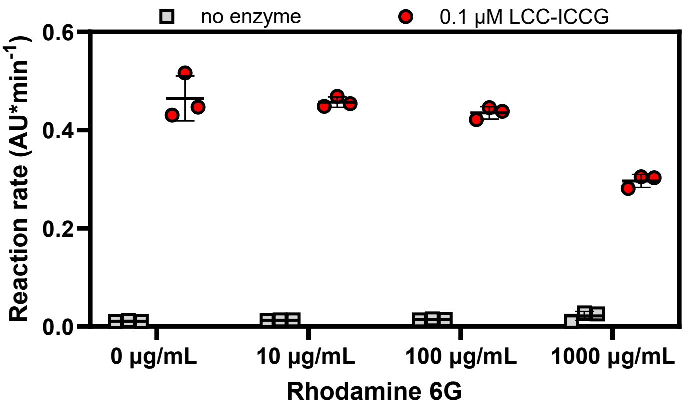


**Figure S4. Effect of rhodamine 6G on the hydrolysis of 4-nitrophenyl acetate by LCC-ICCG.** The reactions were carried out at room temperature in 50 mM Tris-HCl, pH 8.0, using 1 mM substrate and 0.1 µM enzyme. Error bars indicate standard deviations between triplicate measurements (all individual data is shown), whereas horizontal lines denote average values. Reaction progress was followed by monitoring the absorbance of 4-nitrophenol at 400 nm. The reaction rates were determined using the linear (initial) parts of the progress curves. Source data are provided as a Source Data file.

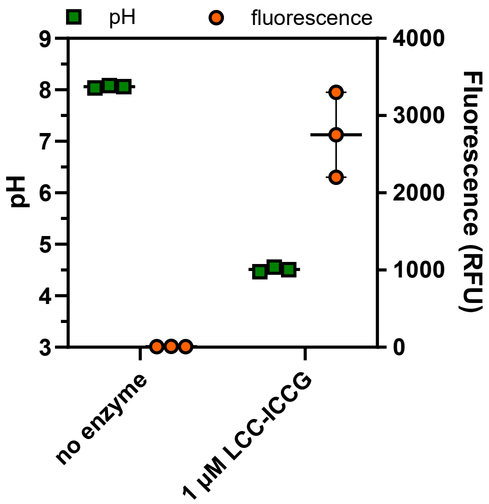


**Figure S5. Acidification of reaction mixtures during enzymatic hydrolysis of PET-R6G.** The reactions were carried out for 96 h at 65 °C in 50 mM Tris-HCl, pH 8.0 using one PET-R6G pellet (~17 mg; 1.7% w/v solids loading) and 1 μM (28.8 μg) LCC-ICCG (1.7 mg enzyme/g PET). Fluorescence was recorded after diluting the sample ten times with the reaction buffer. Error bars indicate standard deviations between replicates (*n* = 3) whereas horizontal lines denote average values. Source data are provided as a Source Data file.

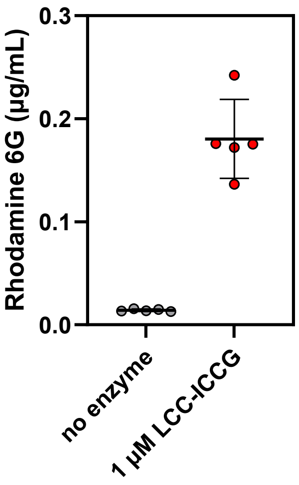


**Figure S6. Enzymatic hydrolysis of PET-R6G at 37 °C.** The 1 mL reaction mixtures containing 50 mM Tris-HCl buffer, pH 8.0, one substrate pellet (≈17 mg; 1.7% w/v solids loading) and 1 μM (28.8 μg) LCC-ICCG (1.7 mg enzyme/g PET) were incubated at 37 °C, 500 RPM for 19 h. 200 μL aliquots were taken from reaction mixtures and transferred to a 96-well microtiter plate. Fluorescence was measured immediately ($\lambda_{ex/em}$ = 530/552 nm) and the concentration of rhodamine 6G was determined according to a standard curve. Error bars indicate standard deviations between replicate measurements (*n = 5*) whereas horizontal lines denote average values. All individual data points are shown. Source data are provided as a Source Data file.

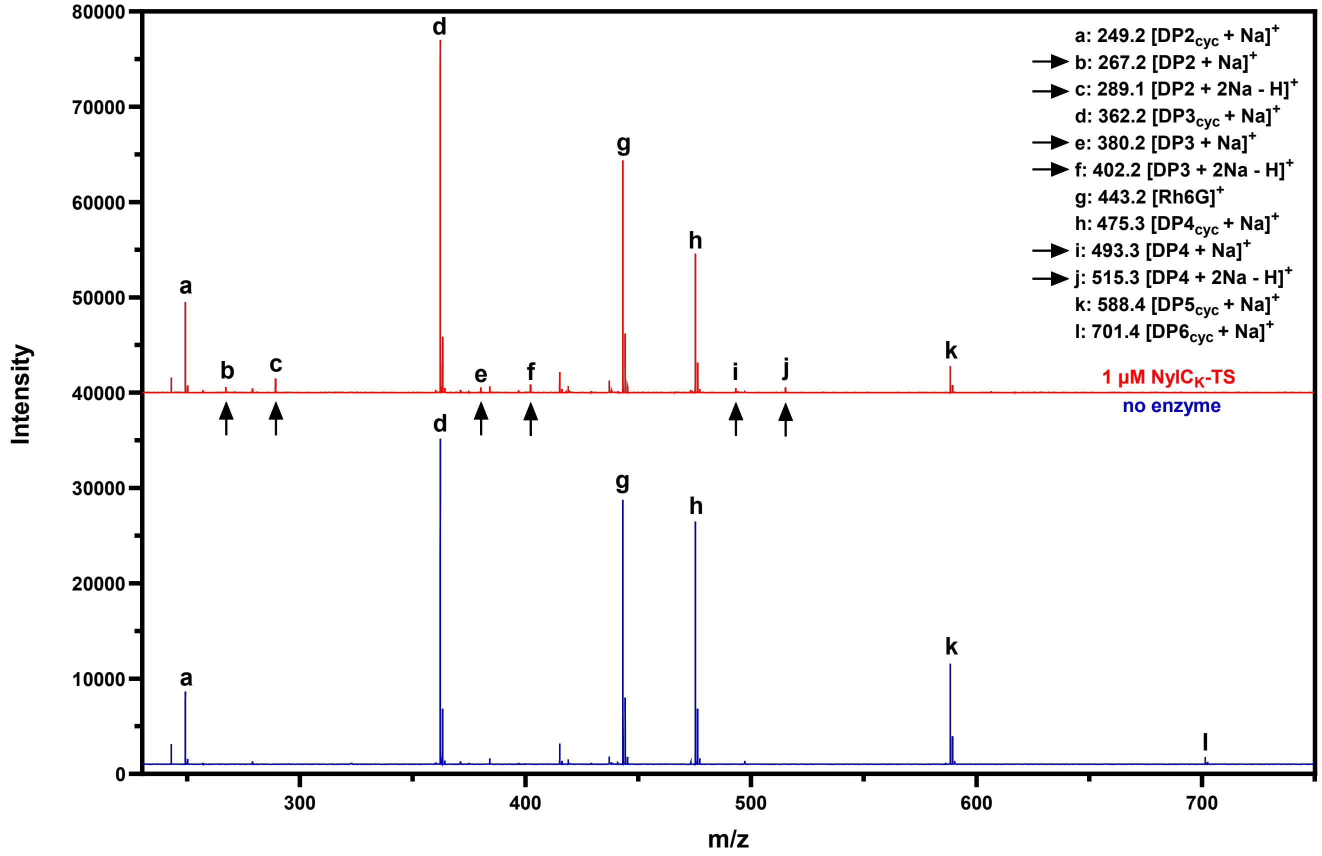


**Figure S7. NylC$_K$-TS activity on PA6-R6G.** The MALDI-ToF MS spectra were obtained after 265 hours of incubation of PA6-R6G (~11 mg; 1.1 % w/v solids loading) in the presence or absence of 1 µM (39 µg; 3.5 mg enzyme/g PET) NylC$_K$-TS in 1 ml of 50 mM sodium phosphate buffer, pH 7.4, supplied with 150 mM NaCl. The experiments were carried out at 65°C, 500 RPM. Cyclic and linear oligomers of nylon-6 were observed as sodium adducts, whereas rhodamine 6G was detected as a free cation. The linear products released upon enzyme addition are marked with arrows. Note that cyclic oligomers with a degree of polymerization of 2-6 ("DP2-6$_{cyc}$") were present in the control reaction lacking the nylonase, indicating substrate contamination (see main text for more discussion). The mass spectra indicate depletion of cyclic hexamer ("l") and cyclic pentamer ("k") in the reaction with enzyme, suggesting that some of the linear products were generated as a result of nylonase acting on these contaminants rather than on the polymeric nylon-6 substrate. Source data are provided as a Source Data file.
